# Neuronal correlates of time integration into memories

**DOI:** 10.1101/2023.09.12.557375

**Authors:** Felix Frantzmann, Marius Lamberty, Laurin Braune, Genevieve M. Auger, Nitin Singh Chouhan, Tobias Langenhan, Mareike Selcho, Dennis Pauls

**Author notes:** Correspondence: Dennis Pauls, Department of Animal Physiology, Institute of Biology, Faculty of Life Sciences, Leipzig University, Germany; Mareike Selcho, Department of Animal Physiology, Institute of Biology, Faculty of Life Sciences, Leipzig University, Germany. **Conflict of interest:** The authors declare no competing financial interests.

## Abstract

The circadian clock affects a wide range of physiological processes. Of particular interest is the influence of the clock on memory performance, as circadian dysfunction is associated with age- and disease-related decline in memory. In various species it has been shown that memory performance is regulated by the circadian clock. However, the anatomical and functional connection of the circadian clock and memory neurons has not been described in detail so far. This study now identifies that Diuretic hormone 31 (DH31)-positive clock neurons of the DN1p cluster regulate memory performance. DH31, a functional homolog of the mammalian calcitonin gene-related peptide, plays a crucial role in this process as a clock communication signal. DH31 facilitates memory performance during the night via indirect signalling, while DH31 signals directly to the mushroom bodies restricting memory performance specifically in the evening. This pleiotropic action of DH31 suggests that the circadian clock confines memory performance to a physiological dynamic range.

## INTRODUCTION

Essential information can be encoded as lasting physical changes in the underlying neuronal circuits. Only if information is precisely stored and retrieved a memory can deliver expected benefits. Therefore, it is not surprising that circadian rhythms, that regulate various physiological processes throughout phylogeny, also have an impact on memory performance (*1–4*). Time information from the circadian clock allows an individual to be in the right place at the right time, which increases the probability of the expected benefit due to the encoded information. It is well established that both invertebrate and vertebrate species display time of day-specific regulation of memory performance, surprisingly, however, we do not know much about the anatomical correlates of time integration in memories. In the last decades, a significant progress has been made in the understanding of the genetic and molecular processes of clock cells. In humans, pacemaker cells in the suprachiasmatic nucleus (SCN) receive light information as a main Zeitgeber signal through the retinohypothalamic tract and generate circadian rhythms, which in turn regulate various physiological processes accordingly (*3*). Importantly, an anatomical connection from the circadian clock to memory-related structures as the hippocampus and striatum is suggested through the paraventricular thalamic nucleus (*5*). Whether these connections are indeed functionally relevant for the time of day-dependent regulation of memory performance is, however, not yet shown unambiguously.

Associative olfactory memories are regulated by the circadian clock also in *Drosophila* (*2*, *6*, *7*). In detail, flies which are trained to form associative olfactory memories show significant learning throughout the day with only modest fluctuations in performance. In contrast, arrhythmic flies lacking the core clock gene *period* or *timeless* do not display any rhythm in memory performance, indicating that the regulation is based on a functional circadian clock (*2*, *6*). This raises the question of how the circadian clock integrates the time of day into neuronal networks underlying memory formation and recall. No expression of clock genes was shown in mushroom body (MB) neurons, so that the rhythm must be generated extrinsically (*8*). A current hypothesis is that the time of day can be encoded by the activity of different clock neurons. The *Drosophila* clock network consists of around 150 neurons in which the conserved core clock machinery generates molecular circadian rhythms. Clock neurons can be distinguished into dorsal neurons (DNs) and lateral neurons (LNs) according to their cell body position in the central brain (*9*). LNs and DNs show a characteristic diurnal activity pattern (*10*), so signals from specific clock neurons at the specific time of day in the MB could provide the temporal context. In this respect, Flyer-Adams et al. provide evidence, that loss of the neuropeptide Pigment-Dispersing-Factor [PDF; (*11*)] reduces learning performance throughout the day, but more substantially in the morning than in the evening (*7*). This corresponds to the activity pattern of the PDF neurons, since neuronal activity is strongest in the morning (*10*). Interestingly, PDF-positive sLNv pacemaker neurons in *Drosophila* show circadian changes in spontaneous firing rate and resting membrane potential like pacemaker cells in the mammalian SCN. This may support the idea of highly conserved neurophysiological mechanisms between the clock and memory circuits (*3*, *12–16*). However, functional imaging data (*7*) suggest that there is no direct PDF signalling to MB intrinsic neurons, i.e. the Kenyon cells (KCs), although the PDF receptor is expressed in KCs according to transcriptomics data (*17*). Therefore, it remains unclear how time information is integrated into memories or how memories can be regulated daytime-specifically.

In the current study, we therefore concentrated on PDFR-positive DN1p neurons (*18*), a cluster of dorso-posterior clock neurons with arborizations in the dorsal protocerebrum. DN1p neurons appear to be non-autonomously and dependent on PDF signalling for clock function (*19–23*). Thus, DN1p neurons appear to be good candidates to convey clock-dependent time information. We here confirm previous findings showing that the circadian clock modulates associative odour-shock memory performance in *Drosophila*. For the first time we provide evidence of a direct anatomical and functional connection between DN1p clock neurons and MB KCs. We show that the lack of Diuretic hormone 31 (DH31), a neuropeptide homolog of vertebrate Calcitonin gene-related peptide and its cognate G protein-coupled receptor (GPCR) DH31R, results in reduced and more variable memory performance. We also suggest that DH31 is not only required for memory facilitation, but also necessary to restrict memory performance during the subjective evening as the specific knockdown of DH31R in the mushroom bodies enhances learning scores. This indicates that the circadian clock homeostatically confines associative learning to a physiological dynamic range.

## RESULTS

### Daytime-specific regulation of memory performance depends on DN1p neurons

Previous work has demonstrated that PDF neurons are required for stable memory performance throughout 24h in *Drosophila*. Interestingly, the MB does not seem to be a direct target of PDF signalling in this context (*7*). Motivated by this finding, we turned our attention to DN1p neurons, a group of 15 clock neurons per side located in the dorsal protocerebrum driving both morning and evening anticipation (*22*, *24*). Importantly, cAMP and calcium are increased in DN1p neurons in response to PDF application. PDF acutely depolarizes and increases action potential firing rates of DN1p neurons (*25*, *26*), proving that DN1p neurons are direct downstream targets of PDF signalling (*24*, *27–29*). To anatomically confirm these findings of overlap between PDF and DN1p neurons, we used the GRASP technique which allows to label potential synaptic contacts by reconstituted GFP fluorescence (*30*). GFP signals became exclusively visible in the superior lateral protocerebrum [Fig.1A; (*26*)]. Analysis of EM data of this particular region using FlyWire (flywire.ai) provided further evidence that DN1p neurons are connected to PDF releasing neurons of the sLNv cluster [Fig.1B- B’’’; (*31*)]. In detail, EM data illustrates dense core vesicles (DCVs) and release sites of a sLNv neuron, as described by (*32*), in close vicinity to a DN1p neuron (arrows; Fig.1B’-B’’’). DN1p neurons are not only downstream of PDF signaling, but also show strong arborizations in the posterior dorsal protocerebrum alongside the MB calyces (Fig.1C-E’). In detail, we found innervation in the region of the dorsal accessory calyx (dAC) which receives rather visual than olfactory input [Fig.1E-E’; (*33*, *34*)]. Thus, it is tempting to speculate that DN1p neurons are required to convey time information from PDF-positive sLNv neurons to the MBs.

**Figure 1:**
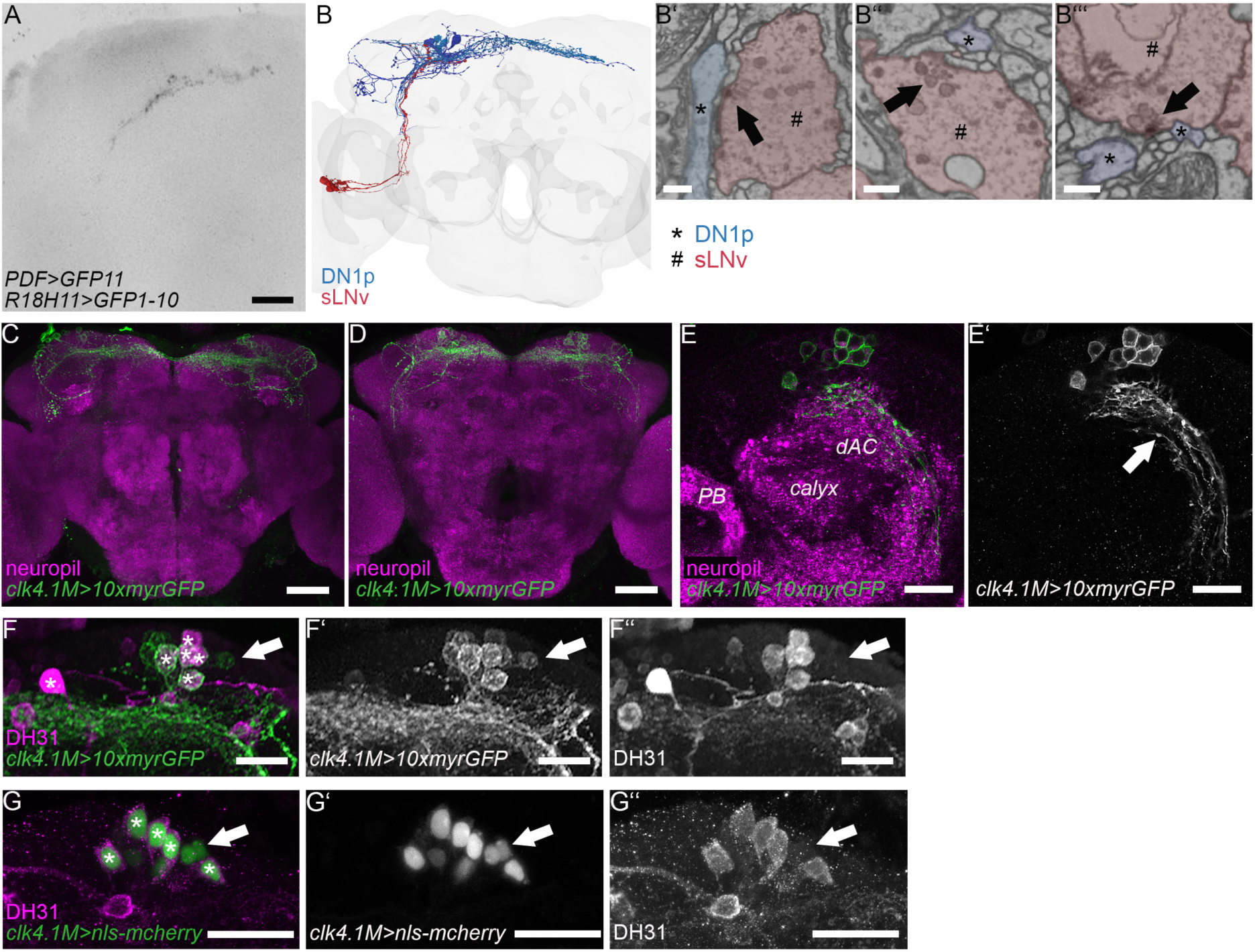
DN1p neurons as prime candidates to convey time information to the MBs. **(A)** Reconstituted split-GFP between PDF-positive sLNvs and DN1p neurons is only visible in the dorsal protocerebrum. **(B-B’’’)** PDF-positive sLNv neurons (red) are connected to DN1p neurons (blue) indicated by dense core vesicle release sites (black arrows; # marks sLNv neurons, * marks DN1p neurons). **(C-E)** Whole mount projections of clk4.1M>10xmyrGFP fly brains (C: anterior view, D: posterior view). DN1p neurons arborize in the dorsal posterior protocerebrum. **(E-E’)** Projection of sections including the dorsal accessor calyx (dAC). DN1p neurons innervate the region of the dAC (arrow) allowing to speculate that they convey time information during memory processes. **(F-G’’)** A subset of DN1p neurons expresses DH31 (*). Green: clk4.1M>10xmyrGFP in (F) and clk4.1M>nls-mcherry in (G), magenta: DH31. [dAC: dorsal accessory calyx; DH31: Diuretic hormone 31; DN1p: dorsal posterior neuron; PB: protocerebral bridge; sLNv: small ventral lateral neuron]

DN1p neurons express DH31 (*18*, *35*) and various other neurotransmitter like glutamate (*22*, *36*), allatostatin-C (*37*), and IPNa (*38*). Interestingly, a hierarchical regulation of rhythmic locomotor behaviour was described for PDF cells and DH31- positive DN1p neurons (*39*). Therefore, we started investigating the role of DH31 in the daytime-specific regulation of memory performance.

DH31 is broadly expressed (Fig.S1A,B), but specifically in a subset of DN1p clock neurons [(*18*, *23*); Fig.1F-G’’]. To challenge the idea that DH31 is essential for stable memory performance throughout 24h found in wildtype flies (Fig.S2), we tested whether a loss of DH31 affects odour-shock learning. We decided to test flies in the evening (ZT9-11), in the early night (ZT13-15) and later during the night (ZT17-19) because normalized memory performance in *pdf^01^* mutant flies was most variable at these times (*7*). In addition, it has been shown that cellular activity in DH31-positive DN1s peaks during the night (*10*). DH31 peptide mutant flies showed reduced memory performance at ZT9-11 and ZT17-19 (Fig.2A,C). On the contrary, mutant flies performed indistinguishable from controls at ZT13-15 likely indicating a day-time specific effect on memory performance (Fig.2B). As these results were based on a null mutation in the DH31 gene (Fig.S3A), we next decided to monitor the flieś circadian activity. In case a mutation of DH31 generally disrupts the function of the circadian clock completely, a statement on a daytime-specific modulation of learning processes would not be possible. In line with previous reports (*39*), the loss of DH31 had no remarkable effect on the flieś rhythmicity (proportion of rhythmicity: control flies 87%, DH31^KG09001^ 93%), at least under LD conditions (Fig.S4A,B). Next, we addressed whether DH31-positive DN1p or LPN neurons, respectively, are required for the effect on memory performance. A recent study has shown that DH31 is also expressed in lateral posterior clock neurons (LPN) with an expression peak in the morning and evening. Interestingly, LPNs were shown to be synaptically connected to MB KCs (*35*). Thus, we used a specific knockdown of *amontillado* in DN1p (via *clk4.1M-Gal4*) and LPN (via *R11B03-p65.AD;R65D05-DBD*) neurons. *Amontillado* encodes the pro- protein convertase dPC2, an enzyme required for neuro- and enteroendocrine peptide synthesis (*40*, *41*). The lack of peptide processing in DN1p neurons reduced memory performance in *clk4.1M>amon-RNAi* flies compared to genetic controls at both ZT9-11 and ZT17-19 (Fig.2D,F). Conveniently, memory performance appeared normal again at ZT13-15 resembling the results obtained earlier as the lack of DH31 affected only ZT9-11 and ZT17-19 (Fig.2A,C). This underlines the idea that the DH31-dependent effect on memory performance is a specific circadian effect.

**Figure 2:**
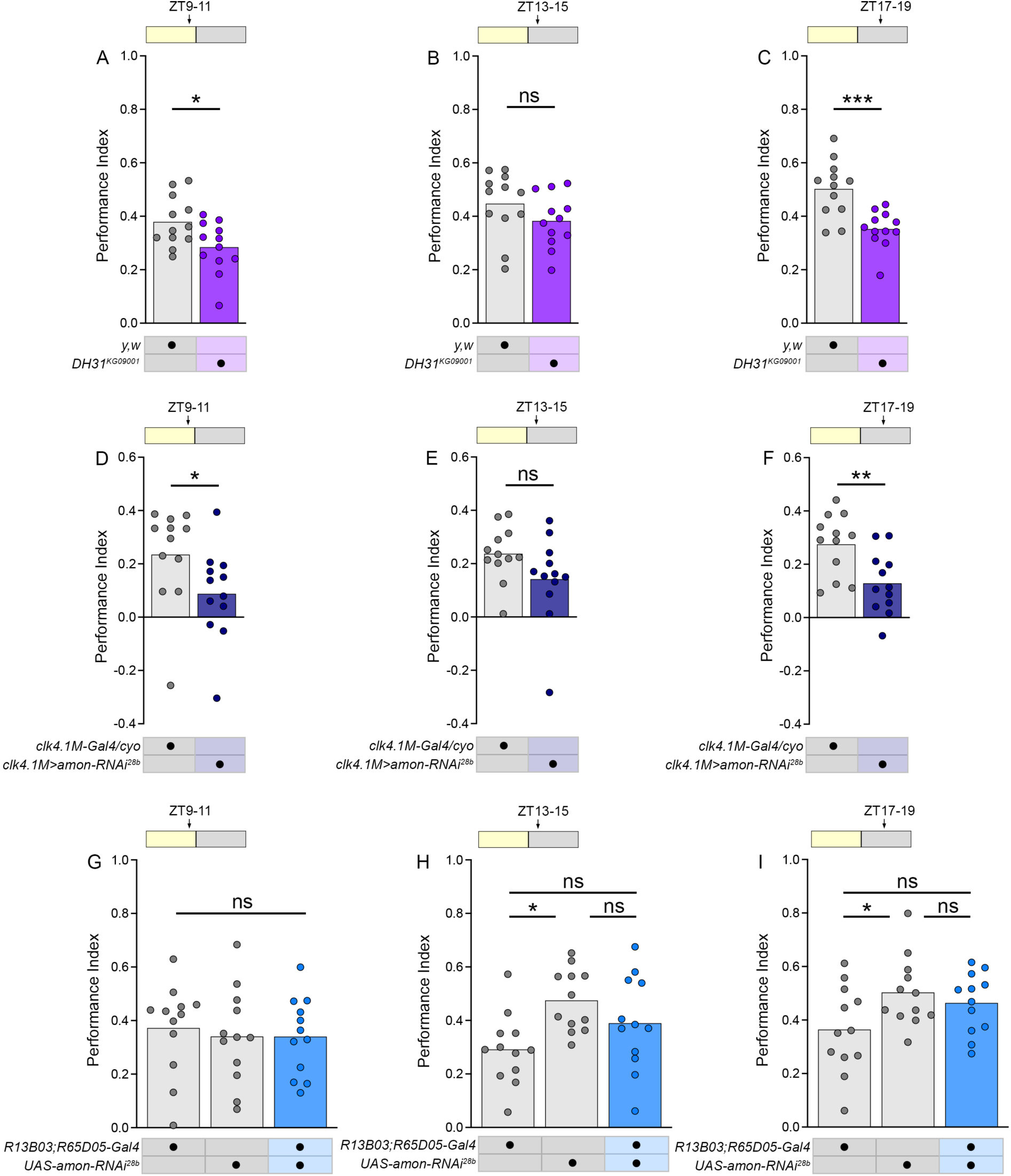
DN1p neurons are involved in the circadian modulation of associative odour-shock memories. **(A-C)** DH31 mutant flies shows impaired learning at ZT9-11 **(A)** and ZT17-19 **(C)**, while they perform indistinguishable from control flies at ZT13-15 **(B)**. **(D-F)** Similarly, the specific knockdown of amontillado in DN1p neurons (via clk4.1m-Gal4) leads to significantly reduced learning scores in the evening [ZT9-11; **(D)**] and at midnight [ZT17-19; **(F)**], while memory performance appears to be normal at ZT13-15 **(E)**. **(G-I)** In contrast, the specific reduction of amontillado in LPN neurons does not affect memory performance. [Amon: amontillado; DH31: Diuretic hormone 31; DN1p: Dorsal posterior neuron; LPN: Lateral posterior neuron; ZT: Zeitgeber time]

The lack of peptide processing in LPN neurons (Fig.2A,B), however, did not affect memory performance at all, indicating that the time-specific and DH31-dependent modulation of memory performance is detached from LPN cell function, at least for the time slots tested. Thus, interfering with DH31 signalling resulted in a reduced memory performance at specific times supporting the idea that DN1p neurons are important to convey time information during memory processes.

### DH31-positive DN1p are directly connected to the mushroom bodies

Our data so far provided evidence that DH31 signalling is required for time-of-day specific regulation of memory performance and that DH31 derives from DN1p neurons.

To confirm a direct modulation of MB KCs through DH31, we analysed the whole brain EM volume using Flywire (*31*, *42*, *43*) and found DCV release sites in a type B DN1p neuron, which is cryptochrome- and DH31-positive (Fig.S1C-1D), next to a ⍺/ß_p_-type KC (asterisks; Fig.3A).

**Figure 3:**
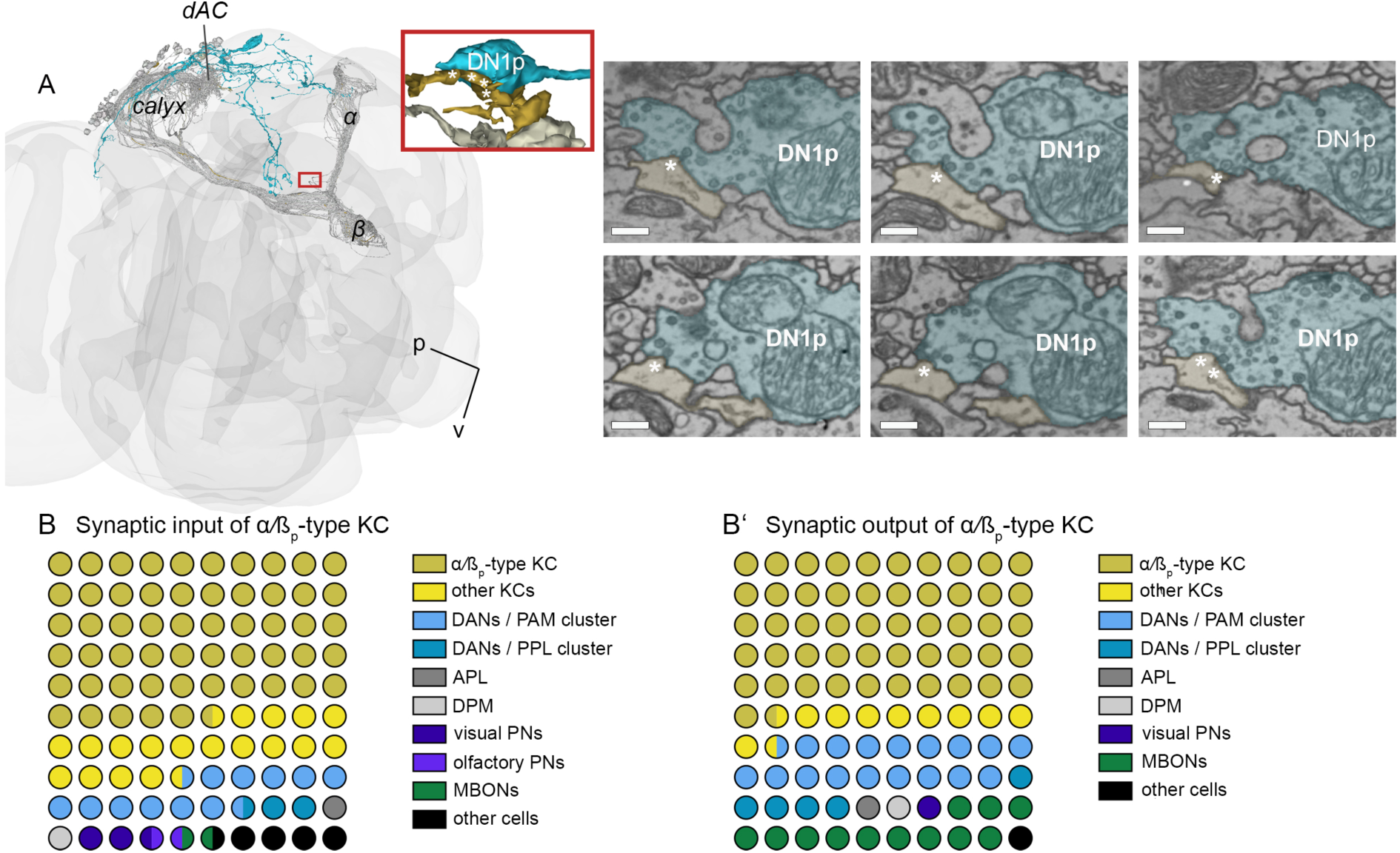
DN1p clock neurons are directly connected to memory neurons in the accessory calyx of the mushroom bodies. **(A)** DCV release sites are visible in a type B DN1p neuron (light blue) connected to an ⍺/ßp-type KC (light yellow). The red square visualizes the contact site. The EM pictures show potential dense core vesicle release sites at the membrane of the DN1p cell next to the KC (asterisks). **(B-B’)** Synaptic input and output connectivity of the target ⍺/ßp-type KC shown in a 10×10 dot plot. The 10×10 dot plot displays the number of synaptic connections found as parts of whole and not exact numbers. [APL: Anterior paired lateral (neuron); dAC: Dorsal accessory calyx; DAN: Dopaminergic neuron; DH31R: Diuretic hormone 31 receptor; DN1p: Dorsal posterior neuron; DPM: Dorsal paired medial (neuron); KC: Kenyon cell; MB: Mushroom body; MBON: Mushroom body output neuron; P: posterior; PAM: Protocerebral anterior medial (cluster); PN: Projection neuron; PPL: Protocerebral anterior lateral (cluster); V: Ventral; ZT: Zeitgeber time]

To provide a further understanding of the corresponding circuit, we characterized synaptic in- and output partners of the identified ⍺/ß_p_-type KC (*44*, *45*). ⍺/ß_p_-type KCs are shown to arborize in the ⍺/ß-lobes while their dendrites extend to the dAC, which is located dorsolateral to the main calyx (*33*, *34*). We identified that the synaptic input comes mainly from other ⍺/ß_p_-type KC. In addition, synaptic input is provided by ⍺/ß_s_- and ⍺’/ß’_m_-type KCs, dopaminergic neurons of the PAM and PPL cluster, projection neurons, the DPM and APL neuron as well as MB output neurons [MBONs; (Fig.3B)]. Interestingly, the composition of synaptic downstream partners was found to be basically the same cell types (Fig.3B’). To challenge the connection between DN1p neurons and KCs functionally, we next specifically downregulated DH31R in MB KCs using RNAi. For this purpose, we chose to use *MB247-Gal4* as this line was shown to drive expression in ⍺/ß_p,s,c_-type KCs and ɣ_d,main-_type KCs, while no expression is visible in ⍺’/ß’-type KC (*46*). This fits RNA-seq data showing that DH31R expression is detectable mainly in ⍺/ß_p,c_-type KCs (*17*). Compared to genetic controls, *MB247>DH31R-RNAi* flies showed significantly increased learning in the evening (ZT9-11; Fig.4A). In contrast, memory performance of *MB247>DH31R-RNAi* flies was indistinguishable from genetic controls at ZT13-15 and ZT17-19 (Fig.4B,C). The increase in memory performance seems to contradict the general reduction in memory performance that occurred with impaired DH31 peptide signalling (Fig.2A-F). To verify the results, we tested memory performance in DH31R mutant flies (Fig.4D-F, FigS2,S3B,S4C). DH31R mutant flies indeed showed reduced memory performance at ZT13-15 and ZT17-19 (Fig.4E,F), which is in line with our results using DH31 peptide mutant flies (Fig.2). In contrast, however, DH31R mutant flies were indistinguishable from wildtype flies at ZT9-11 (Fig.4D), which appears to be in line with the specific knockdown of DH31R in MB KCs and the corresponding increase in memory performance. Taken together, our data suggests that odour-shock learning is confined to a dynamic range by the circadian clock. Here, DH31 signalling seems to facilitate memory performance at certain times of the day, while direct DH31:MB signalling seems to be necessary specifically for the restriction of memory performance in the evening.

**Figure 4:**
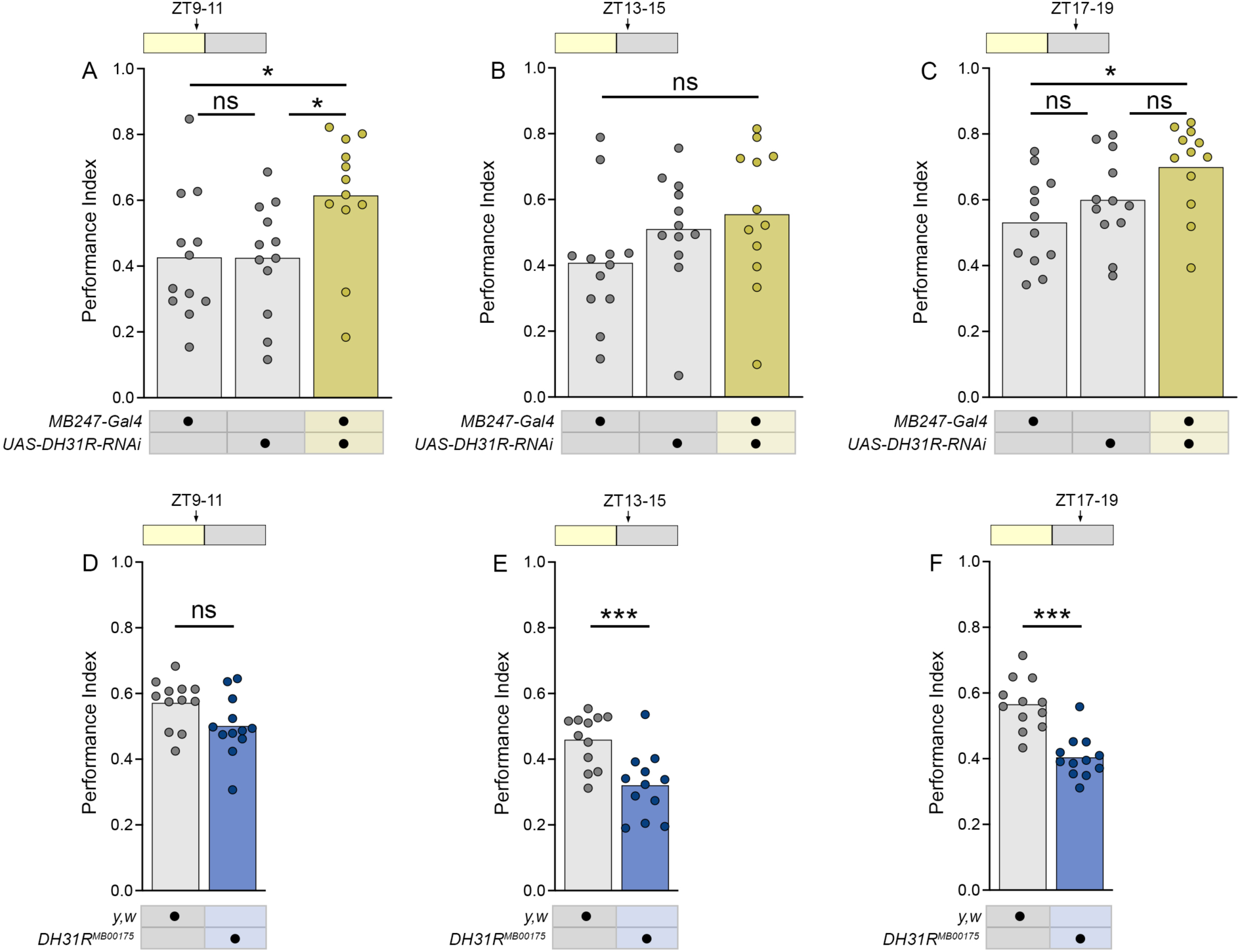
Associative memory performance is restricted in the subjective evening through direct DH31 signalling. **(A-C)** Specific knockdown of DH31R in MB KCs (using MB247-Gal4) resulted in increased learning scores at ZT9-11 **(A)**, while learning performance was indistinguishable from genetic controls at ZT13-15 **(B)** and ZT17-19 **(C)**. **(D-E)** Memory performance of DH31R mutant and control flies. DH31R mutant flies showed significantly reduced at ZT13-15 **(E)** and ZT17-19 **(F)** compared to control flies. At ZT9-11, however, DH31R mutant flies performed indistinguishable compared to controls indicating an elevated or rather non-reduced memory performance in the evening **(D)**. [ZT: Zeitgeber time]

## DISCUSSION

A circadian dysfunction is coupled with adverse fitness consequences, sleep disorders, metabolic and cardiovascular diseases, but also with age-related and disease-related decline in memory including retrograde amnesia (*47–51*). Neurodegenerative diseases such as Alzheimeŕs disease and Parkinson disease as well are often observed to be interrelated with circadian system deterioration (*52*). Thus, the question arises how the circadian clock impacts on memory acquisition and/or memory recall. Previous studies have already shown that both vertebrate and invertebrate species are able to specifically integrate the time of day into long-term memory processes . Here, memory performance is not modulated, but rather tagged with a specific time-stamp.

Accordingly, memory retention scores appeared to be significantly higher when rats were tested either immediately after training or at successive multiples of 12h later than tested 6h after training (*56*, *57*). Likewise, flies can learn to specifically approach an odour that was associated with reward in the morning, while they approach a second odour, that was coupled with the same reward in the afternoon, at the corresponding time of day (*58*, *59*). Importantly, constant light interferes with the flieś ability for time- odour learning, emphasizing the importance of a functional circadian clock.

However, there is growing evidence that the circadian clock also regulates short-term memory performance throughout 24h without mapping a specific time stimulus onto CS-US associations. Various species show peak performance of memory at most active times, such as zebrafish (*D. rerio*) and the sea slug (*A. californica*), while others peak in memory performance in their inactive phase such as mice (*M. musculus)* and flies (*D. melanogaster*) (*1–3*, *60–62*). Notably, the mild peak in memory performance was not visible in our study, which could be due to different methods used in entrainment. Our study substantiates that memory performance is regulated by the circadian clock. As odour memories are stored in the MBs, the question is whether KCs can self-autonomously integrate time information or whether the circadian clock must convey the information during conditioning. 1144 genes were identified to be expressed rhythmically in MB KCs. However, none of these genes were shown to be canonical clock genes indicating that rhythmic gene expression in these memory neurons must be controlled extrinsically through clock function (*63–65*). Surprisingly, a direct, functional connection of the clock and the MBs was not shown so far. sLNvs arborize dorsofrontally of the MB calyces (*32*) providing a potential neuronal link between the clock and circuits associated with memory formation (*66–69*). However, although PDFR mRNA expression was found in subpopulations of KCs, they might not be direct functional targets of PDF signalling (*7*, *17*). Nevertheless, lack of PDF/PDFR signalling impaired appetitive STM expression suggesting that these core pacemaker cells are indeed required for normal memory performance throughout 24h (*7*). We now found similar results focusing on DH31. DH31 is a functional homolog of the mammalian calcitonin gene-related peptide (CGRP) and has been shown to be a circadian wake-promoting signal in flies (*18*). Interestingly, CGRP immunoreaction was found in the SCN of the mouse hypothalamus and in cells of the intergeniculate leaflet (IGL) of the thalamus. While the SCN is known to harbour main pacemaker cells generating circadian rhythms in mammals, cells of the IGL send arborizations to the SCN. By mediating light information IGL cells have a strong impact on circadian rhythms (*70*, *71*). The corresponding CGRP receptor complex consists of a class B GPCR (calcitonin receptor-like receptor, CLR), a receptor activity modifying protein 1 (RAMP) and a receptor component protein (RCP) (*72*). While circadian dysfunction in Alzheimeŕs disease patients appear to be more severe, CGRP appears to be a target site for treatment which further underlines the importance of the circadian clock on memory performance and pathophysiology and the importance of CGRP/DH31 as a communication signal (*73*).

Flies lacking DH31/DH31R signalling display impaired and more variable STM performance. Our data suggests that DH31/DH31R signalling downstream of PDF signalling is required for conveying time information indirectly to the MBs as the specific knockdown of DH31R in MB KCs did not reproduce the learning deficit from the mutant flies at ZT13-15 and ZT17-19. However, it is well possible that this functional difference is due to diurnally regulated variations in the arborization pattern of DN1p in the MB circuitry. The connectomics analysis in this study identified a few distinct points of contact between DN1ps and KCs, revealing for the first time the direct connection of the clock to the memory centre. It is now easy to imagine that these arborizations and contact points increase significantly at night, for example, as at this time DN1ps peak in activity (*10*). Such a diurnal adjustment of the arborizations was also shown for the PDF-positive sLNvs (*74*, *75*) and in DN1a neurons (*76*). Interestingly, DH31 signalling appeared to be not exclusively important for the facilitation of memories, but also for the limitation at ZT9-11. Here, the specific knockdown of DH31R in MB KCs resulted in an enhanced memory performance indicating pleiotropic action of DH31 on memory performance through an indirect and direct signalling to the MBs, respectively. Another explanation is a more complex circuit in which DH31R-positive neurons receive further afferents, so that DH31 signals are not absolutely essential for the activity of DH31R- positive neurons. Here, it is of particular interest that our connectomics analysis revealed DH31R-positive KCs to be embedded in a well-known complex network including the APL, DPM, MBINs and MBONs allowing different feedback and feedforward signalling pathways (*41*, *67*, *68*, *77–79*).

A circadian regulation of memory performance is important to maintain learning capabilities constant and to adapt to physiological or environmental changes. This allows the organism to limit the unnecessary depletion of energy and avoid any fitness consequences. Thus, it appears coherent that learning is impaired through a disrupted circadian clock. At such a state, it is not possible to maintain energy-demanding memory processes to the full extent. Indeed, our study confirms previous work that impairing the circadian clock results in a rather general decrease of learning performance in flies (*7*). However, a common task of the circadian clock may also include the restriction of learning. Zebrafish perform better in an operant learning task during the day than at night. Here, clock-derived melatonin appeared to be both necessary and sufficient for the reduced memory scores during the night (*61*). Although learning scores were constant throughout 24h in our study in control flies, our data may resemble the inhibitory function of the circadian clock on learning and memory. First, control flies showed a slight but significantly reduced learning in the early night (ZT 13- 15) compared to the early day (ZT 1-3). Second, specific impairment of DH31 signalling to the MBs increased learning scores in the evening, suggesting such an inhibitory function of the circadian clock on learning and memory. It is tempting to speculate that the limitation of learning capacity may rely on metabolic energy regulation. Brain functions, in particular memory acquisition and consolidation, are highly energy demanding and thus fine regulations of the energy metabolism are crucial (*80*). Indeed, appetitive odour-sugar memories in *Drosophila* are suppressed by satiety while they are promoted by hunger (*81*). Furthermore, food deprived flies defer from forming energy demanding aversive long-term memory in favour of a low-cost anaesthesia-resistant memory, which is independent on de novo protein synthesis (*82*). In the same way, starvation is essential for the acquisition, but not for the retrieval of time-odour memories (*59*). Interestingly, extra food deprivation is required in female flies to achieve a similar time-dependent memory performance as compared to male flies, which is probably due to different energy demands between male and female flies (*59*). Thus, it is hypothesized that one specific function of the circadian clock is the state-dependent limitation of learning processes. Flies display a clock-dependent feeding rhythm with a major peak in the early morning with most moderate temperature and humidity conditions, which mutually competes with other behaviours like sleep (*83*, *84*). Limiting learning performance, for example in the evening as shown in the present study, would therefore allow the regulation of energy supplies rationally.

## Acknowledgements

The authors thank Robert J. Kittel, Andreas S. Thum, Charlotte Helfrich-Förster and Martin Heisenberg for fruitful discussions and/or support provided by equipment, Konrad Öchsner, Hans Kaderschabek, and Nicole Naumann for excellent technical assistance, Michael Nitabach, Jan Veenstra and Pamela Menegazzi for providing antibodies, Dirk Rieger and the Bloomington Stock Center for providing flies. We thank the Princeton FlyWire team and members of the Murthy and Seung labs, as well as members of the Allen Institute for Brain Science, for development and maintenance of FlyWire (supported by BRAIN Initiative grants MH117815 and NS126935 to Murthy and Seung). We also acknowledge members of the Princeton FlyWire team and the FlyWire consortium for neuron proofreading and annotation. We thank members of the Dickson, Murthy, Seung, Kim, and Jeffries lab for proofreading and annotations of the cells shown in this publication (Table S1). This work was supported by the Deutsche Forschungsgemeinschaft (DP1979/2-1, DP1979/3-1 and DP1979/5-1 to D.P., PA3241/2-1 to M.S., through CRC 1423, project number 421152132 [project B06] to T.L.) and a PhD fellowship (to F.F.) from the German Excellence Initiative to the Graduate School of Life Sciences, University of Würzburg, Germany. The authors declare no competing interests. F.F., L.B., N.C., T.L., M.S. and D.P. conceived and designed the experiments. F.F., M.L., L.B., G.A., M.S. and D.P. performed the experiments. F.F., L.B., M.S. and D.P. analysed the results. F.F., M.S., and D.P. wrote the article. All authors provided comments and approved the manuscript.

## METHODS

### Fly strains

All fly strains (Tab.1) were reared in vials containing standard cornmeal medium at 25 °C and 65 % relative humidity with a 12:12 LD cycle.

**Table 1:** Fly lines used in this study.

| Driver lines | Genotype | Reference |
| --- | --- | --- |
| *clk4.1m-Gal4* | w; sco/cyo;clk4.1M-Gal4/TM6b | (24) |
| *MB247-Gal4* | w; +, MB247-Gal4 | (85) |
| *R13B03,R65D05-Gal4* | w; R11B03-p65.AD;<br>R65D05-DBD | (86) |
| *R18H11-Gal4* | w[1118];P{y[+t7.7]<br>w[+mC]=GMR18H11-GAL4}attP2 | BDSC #48832 |
| *PDF-lexA* | yw; PDF-lexA; + | (87) |
| Reporter lines | Genotype | Reference |
| UAS-*amon-RNAi*<sup>28b</sup> | w; UAS-*amon-RNAi*28b; + | (88) |
| UAS-*DH31R-RNAi* | w; UAS-dh31r-RNAi; + | VDRC #8777 |
| 10xUAS-IVS-*myr::GFP* | w[*]; P{y[+t7.7] w[+mC]=10XUAS-<br>IVS-*myr::GFP*}attP2 | BDSC #32197 |
| UAS-*nls-mcherry* | w; +; UAS-mCherry.NLS | BDSC #38424 |
| LexAop-spGFP11; UAS-spGFP1-10 | w; LexAop-GFP11/cyo; UAS-GFP1-10/TM6b | (89) |
| Other lines | Genotype | Reference |
| DH31<sup>KG09001</sup> | y; P{SUPor-P}Dh31KG09001 | BDSC #16474 |
| DH31R<sup>MB00175</sup> | y w; Mi{ET1}Dh31-R[MB00175] | BDSC #22718 |
| *w*<sup>1118</sup> | w; +; + | --- |
| Yellow-white | yw; +; + | --- |

### Behavioural assays

#### Olfactory associative conditioning

For odour-shock experiments 3 – 7 days old flies were fed on fresh food vials for up to 24 hours prior to behavioural tests. Male and female flies were trained and tested in mixed groups of approximately 100 – 150 animals. Training and testing of flies were done in a modified version of a T-maze assay in which four independent groups of flies can be simultaneously tested. During the experiment, flies were exposed to an airstream of approximately 750 mL/min carrying odours [4-Methylcyclohexanol (MCH, Sigma-Aldrich) and 3-Octanol (OCT, Sigma-Aldrich), which was diluted 1:10 in paraffin oil (Sigma-Aldrich) from attached odour cups. All experiments were done in either dim red light or in complete darkness during the test period. Further, experiments were performed at 23 °C and 80 % humidity.

During aversive conditioning, flies were exposed to the first odour (CS+) for one minute in the presence of 12 pulses of electric shocks (60 V, 1.3 s; US) inside a tube lined with an electrifiable copper grid. After 45 seconds of fresh air, flies received a second odour (CS-) in absence of any US, followed by an additional 45 seconds of fresh air. Afterwards, flies rested for two minutes. Subsequent testing was done in blank Teflon tubes where flies were allowed to choose between CS+ and CS- for two minutes. To avoid skewed associative learning scores by non-associative odour effects, a reciprocal experimental design was used. Two groups of flies either received odour A with electric shocks and odour B without, or alternatively odour B with electric shocks and odour A without. For both groups a learning index was calculated as follows: number of flies choosing CS+ subtracted by the number of flies choosing CS-, divided by the total amount of flies. Memory performance was calculated as the average of the two reciprocally trained groups.

To investigate the impact of the circadian clock on memory performance, aversive olfactory conditioning experiments were performed throughout the flies’ subjective day and night. We have chosen the following time slots, with ZT0 (ZT: Zeitgeber time) defined as start of the day (lights-on): ZT1-3; ZT5-7; ZT9-11; ZT13-15; ZT17-19 and ZT21-23. Flies trained and tested during their subjective day were taken from the entrainment chamber shortly before experiments. In contrast, flies tested at night were taken out of the entrainment boxes at the end of the day and kept in the dark until the experiment to avoid a light pulse at night.

#### Locomotor activity recording

3-7 days old flies were recorded individually in *Drosophila* activity monitors (DAM, Trikinetics, Waltham, USA) under standard 12:12 light:dark (LD) conditions with light intensities around 100-300lux at 25°C for several days. Flies were kept in small glass tubes with 2% agarose and 4% sugar on one side. Fly activity was defined by the number of infrared light beam crosses per minute. Single fly actograms were compiled using the Fiji plugin Actogram J (*90*), while analysis of rhythmicity and activity levels was done with excel.

#### Immunostainings

3-5d old flies were dissected in ice cold 1x PBS and whole mounts were fixed with 4 % paraformaldehyde for 90 min at room temperature. After fixation and washing with 3x 10 min with 0.3 % PBT (1x PBS with 0.3 % Triton X-100; Sigma Aldrich) at room temperature, specimens were blocked 2h with 5% normal goat serum (NGS) in 0.3 % PBT (1x PBS with 0.3 % Tritonx-100; Sigma Aldrich). Incubation with primary antibodies (Tab.2) in blocking solution overnight was followed by 6×10 min washes in 0.3 % PBT and incubation with secondary antibodies. The next day brains were washed 5×10min in PBT and 1×10min in PBS, subsequently mounted in Vectashield H-1000 (Vector laboratories). Whole mounts were stored at 4 °C until scanning with a confocal light scanning microscope (ZEISS LSM800, ZEISS Microscopy). Confocal image stacks were taken at z-step size of 0.93µm using a LD LCI Plan-Apochromat 25x/0.8 Imm Korr DIC M27 objective or a z-step size of 0.44µm using a 63x/1.4 NA oil DIC M27 objective. Intensity and contrast were adjusted using Image J [version 2.9.0/1.53t] and Adobe Photoshop 2023 (Adobe Systems Software).

**Table 2:** Antibodies used in this study.

| Antibody | Manufacturer | Working dilution |
| --- | --- | --- |
| Rabbit anti-DH31 | kindly provided by Jan Veenstra | 1:500 |
| Guinea-pig anti-DH31 | kindly provided by Michael Nitabach | 1:400 |
| Mouse anti-Synapsin | 3C11, DSHB | 1:50 |
| Mouse anti-Fasciclin II | 1D4, DSHB | 1:50 |
| Goat-anti-rabbit-AlexaFluor488 | A11039 | 1:200 |
| Goat-anti-rabbit-STARred | Abberior | 1:200 |
| Goat-anti-mouse-STARred | Abberior | 1:200 |
| Mouse anti-GFP | Sigma-Aldrich, SAB4200681 | 1:500 |
| Goat-anti-mouse AF488 | Invitrogen, A-11001 | 1:250 |

#### EM reconstruction analysis

The EM data and connectomic analysis was performed in FlyWire [(flywire.ai, (*31*, *42*)] and is based on the EM dataset of a female fly brain from (*43*). We analysed all pre- and postsynaptic partners of the KC (FlyWire ID: 720575940617175413) with at least one synapse (*44*, *45*).

### Genotyping of DH31^KG09001^ and DH31R^MB00175^ mutant lines

Genomic DNA was extracted from adult fly homogenate samples using a Nucleospin Tissue Kit (Machery-Nagel, # 740952.250). PCRs for *DH31* and *DH31R* alleles were conducted using the Taq PCR Master Mix Kit (Qiagen, # 201445) and all primers were synthesized by Microsynth (Balgach, Switzerland) using the following primer pairs: *DH31* mutant (primers used: tl_939F: 5’-gatggaaatatacttaggcagctga-3’ and tl_940R: 5’-accgtatcctccattggattggcta-3’), *DH31R* mutant (primers used: tl_941F: 5’- tttatgcagtctaatttacgcactt-3’ and tl_942R: 5’-attgcgttctcctctggcagtcagc-3’), animals genotyped were homozygous. DNA from *w^1118^* was used as negative control for both reactions and an additional control PCR for the presence of the ***α**-Tubulin* gene (primers used: rk_16F: 5’-gtgaattttccttgtcgcgtg-3’ and rk_17R: 5’- ctccagtctcgctgaagaag-3’) was performed on all genotypes. All PCR reactions were performed with a Biometra TRIO Multi-Block PCR Machine (Analytik Jena, Überlingen Germany). The following PCR cycling conditions for the *DH31* and *DH31R* alleles were used: denaturation of dsDNA ran for 3 minutes at 94 °C, followed by 35 cycles consisting of 30 seconds denaturation at 94°C, 30 seconds annealing at 60 °C and 45 seconds of elongation at 72 °C. This was completed by a final elongation step at 72 °C for an additional 10 minutes. The same parameters were used for the ***α**-Tubulin* PCR amplification except the annealing temperature was 55 °C and a total of 40 cycles were run. The PCR products were visualized via gel-electrophoresis using a 1% agarose gel stained with Ethidium bromide (ROTH, # 2218.2).

### Statistics

The data on learning in this study is shown as positive data, even though it is aversive odour-shock learning. Data was analysed for normal distribution using the Shapiro- Wilk test. For statistical comparison between genotypes, an ordinary one-way ANOVA with post-hoc Tukeýs multiple comparison test was used for normally distributed data, a Kruskal-Wallis test was used for not normally distributed data. Single comparisons were made using the Student’s t-test or Mann Whitney test, depending on whether the data was normally distributed. All statistical analyses and visualizations were done with GraphPad Prism 9 (GraphPad Software, San Diego, California, USA) and Adobe Photoshop 2023.

## REFERENCES

1. R. I. Fernandez, L. C. Lyons, J. Levenson, O. Khabour, A. Eskin, Circadian modulation of long- term sensitization in Aplysia. Proc. Natl. Acad. Sci. U. S. A. 100, 14415–14420 (2003).

2. L. C. Lyons, G. Roman, Circadian modulation of short-term memory in Drosophila. Learn. Mem. Cold Spring Harb. N. 16, 19–27 (2009).

3. J. R. Gerstner, J. C. P. Yin, Circadian rhythms and memory formation. Nat. Rev. Neurosci. 11, 577–588 (2010).

4. C. K. Mulder, M. P. Gerkema, E. A. Van der Zee, Circadian clocks and memory: time-place learning. Front. Mol. Neurosci. 6, 8 (2013).

5. M. M. Moga, R. P. Weis, R. Y. Moore, Efferent projections of the paraventricular thalamic nucleus in the rat. J. Comp. Neurol. 359, 221–238 (1995).

6. R. Fropf, H. Zhou, J. C. P. Yin, The clock gene period differentially regulates sleep and memory in Drosophila. Neurobiol. Learn. Mem. 153, 2–12 (2018).

7. J. G. Flyer-Adams, E. J. Rivera-Rodriguez, J. Yu, J. D. Mardovin, M. L. Reed, L. C. Griffith, Regulation of Olfactory Associative Memory by the Circadian Clock Output Signal Pigment-Dispersing Factor (PDF). J. Neurosci. 40, 9066–9077 (2020).

8. P. Machado Almeida, B. Lago Solis, L. Stickley, A. Feidler, E. Nagoshi, Neurofibromin 1 in mushroom body neurons mediates circadian wake drive through activating cAMP-PKA signaling. Nat. Commun. 12, 5758 (2021).

9. C. Helfrich-Förster, O. T. Shafer, C. Wülbeck, E. Grieshaber, D. Rieger, P. Taghert, Development and morphology of the clock-gene-expressing lateral neurons of Drosophila melanogaster. J. Comp. Neurol. 500, 47–70 (2007).

10. X. Liang, T. E. Holy, P. H. Taghert, Synchronous Drosophila circadian pacemakers display nonsynchronous Ca^2+^ rhythms in vivo. Science. 351, 976–981 (2016).

11. S. C. Renn, J. H. Park, M. Rosbash, J. C. Hall, P. H. Taghert, A pdf neuropeptide gene mutation and ablation of PDF neurons each cause severe abnormalities of behavioral circadian rhythms in Drosophila. Cell. 99, 791–802 (1999).

12. D. J. Green, R. Gillette, Circadian rhythm of firing rate recorded from single cells in the rat suprachiasmatic brain slice. Brain Res. 245, 198–200 (1982).

13. C. M. A. Pennartz, M. T. G. de Jeu, N. P. A. Bos, J. Schaap, A. M. S. Geurtsen, Diurnal modulation of pacemaker potentials and calcium current in the mammalian circadian clock. Nature. 416, 286–290 (2002).

14. S. J. Kuhlman, D. G. McMahon, Rhythmic regulation of membrane potential and potassium current persists in SCN neurons in the absence of environmental input. Eur. J. Neurosci. 20, 1113–1117 (2004).

15. D. Park, L. C. Griffith, Electrophysiological and anatomical characterization of PDF-positive clock neurons in the intact adult Drosophila brain. J. Neurophysiol. 95, 3955–3960 (2006).

16. V. Sheeba, H. Gu, V. K. Sharma, D. K. O’Dowd, T. C. Holmes, Circadian- and light-dependent regulation of resting membrane potential and spontaneous action potential firing of Drosophila circadian pacemaker neurons. J. Neurophysiol. 99, 976–988 (2008).

17. M.-F. M. Shih, F. P. Davis, G. L. Henry, J. Dubnau, Nuclear Transcriptomes of the Seven Neuronal Cell Types That Constitute the Drosophila Mushroom Bodies. G3 Genes Genomes Genet. 9, 81–94 (2019).

18. M. Kunst, M. E. Hughes, D. Raccuglia, M. Felix, M. Li, G. Barnett, J. Duah, M. N. Nitabach, Calcitonin gene-related peptide neurons mediate sleep-specific circadian output in Drosophila. Curr. Biol. CB. 24, 2652–2664 (2014).

19. S. Veleri, C. Brandes, C. Helfrich-Förster, J. C. Hall, R. Stanewsky, A self-sustaining, light- entrainable circadian oscillator in the Drosophila brain. Curr. Biol. CB. 13, 1758–1767 (2003).

20. A. Klarsfeld, S. Malpel, C. Michard-Vanhée, M. Picot, E. Chélot, F. Rouyer, Novel features of cryptochrome-mediated photoreception in the brain circadian clock of Drosophila. J. Neurosci. Off. J. Soc. Neurosci. 24, 1468–1477 (2004).

21. T. Yoshii, C. Wülbeck, H. Sehadova, S. Veleri, D. Bichler, R. Stanewsky, C. Helfrich-Förster, The Neuropeptide Pigment-Dispersing Factor Adjusts Period and Phase of Drosophila’s Clock. J. Neurosci. 29, 2597–2610 (2009).

22. A. Chatterjee, A. Lamaze, J. De, W. Mena, E. Chélot, B. Martin, P. Hardin, S. Kadener, P. Emery, F. Rouyer, Reconfiguration of a Multi-oscillator Network by Light in the Drosophila Circadian Clock. Curr. Biol. CB. 28, 2007–2017.e4 (2018).

23. A. Lamaze, R. Stanewsky, DN1p or the “Fluffy” Cerberus of Clock Outputs. Front. Physiol. 10 (2020), doi:10.3389/fphys.2019.01540.

24. Y. Zhang, Y. Liu, D. Bilodeau-Wentworth, P. E. Hardin, P. Emery, Light and temperature control the contribution of specific DN1 neurons to Drosophila circadian behavior. Curr. Biol. CB. 20, 600–605 (2010).

25. O. T. Shafer, D. J. Kim, R. Dunbar-Yaffe, V. O. Nikolaev, M. J. Lohse, P. H. Taghert, Widespread receptivity to neuropeptide PDF throughout the neuronal circadian clock network of Drosophila revealed by real-time cyclic AMP imaging. Neuron. 58, 223–237 (2008).

26. A. Seluzicki, M. Flourakis, E. Kula-Eversole, L. Zhang, V. Kilman, R. Allada, Dual PDF signaling pathways reset clocks via TIMELESS and acutely excite target neurons to control circadian behavior. PLoS Biol. 12, e1001810 (2014).

27. S. Hyun, Y. Lee, S.-T. Hong, S. Bang, D. Paik, J. Kang, J. Shin, J. Lee, K. Jeon, S. Hwang, E. Bae, J. Kim, Drosophila GPCR Han is a receptor for the circadian clock neuropeptide PDF. Neuron. 48, 267–278 (2005).

28. B. C. Lear, L. Zhang, R. Allada, The neuropeptide PDF acts directly on evening pacemaker neurons to regulate multiple features of circadian behavior. PLoS Biol. 7, e1000154 (2009).

29. O. T. Shafer, G. J. Gutierrez, K. Li, A. Mildenhall, D. Spira, J. Marty, A. A. Lazar, M. de la P. Fernandez, Connectomic analysis of the Drosophila lateral neuron clock cells reveals the synaptic basis of functional pacemaker classes. eLife. 11, e79139 (2022).

30. E. H. Feinberg, M. K. VanHoven, A. Bendesky, G. Wang, R. D. Fetter, K. Shen, C. I. Bargmann, GFP Reconstitution Across Synaptic Partners (GRASP) Defines Cell Contacts and Synapses in Living Nervous Systems. Neuron. 57, 353–363 (2008).

31. S. Dorkenwald, C. E. McKellar, T. Macrina, N. Kemnitz, K. Lee, R. Lu, J. Wu, S. Popovych, E. Mitchell, B. Nehoran, Z. Jia, J. A. Bae, S. Mu, D. Ih, M. Castro, O. Ogedengbe, A. Halageri, K. Kuehner, A. R. Sterling, Z. Ashwood, J. Zung, D. Brittain, F. Collman, C. Schneider-Mizell, C. Jordan, W. Silversmith, C. Baker, D. Deutsch, L. Encarnacion-Rivera, S. Kumar, A. Burke, D. Bland, J. Gager, J. Hebditch, S. Koolman, M. Moore, S. Morejohn, B. Silverman, K. Willie, R. Willie, S.-C. Yu, M. Murthy, H. S. Seung, FlyWire: online community for whole-brain connectomics. Nat. Methods. 19, 119–128 (2022).

32. K. Yasuyama, I. A. Meinertzhagen, Synaptic connections of PDF-immunoreactive lateral neurons projecting to the dorsal protocerebrum of Drosophila melanogaster. J. Comp. Neurol. 518, 292– 304 (2010).

33. Y. Aso, D. Hattori, Y. Yu, R. M. Johnston, N. A. Iyer, T.-T. Ngo, H. Dionne, L. Abbott, R. Axel, H. Tanimoto, G. M. Rubin, The neuronal architecture of the mushroom body provides a logic for associative learning. eLife. 3, e04577 (2014).

34. J. Li, B. D. Mahoney, M. S. Jacob, S. J. C. Caron, Visual Input into the Drosophila melanogaster Mushroom Body. Cell Rep. 32, 108138 (2020).

35. N. Reinhard, E. Bertolini, A. Saito, M. Sekiguchi, T. Yoshii, D. Rieger, C. Helfrich-Förster, The lateral posterior clock neurons of Drosophila melanogaster express three neuropeptides and have multiple connections within the circadian clock network and beyond. J. Comp. Neurol. 530, 1507–1529 (2022).

36. Y. Hamasaka, D. Rieger, M.-L. Parmentier, Y. Grau, C. Helfrich-Förster, D. R. Nässel, Glutamate and its metabotropic receptor in Drosophila clock neuron circuits. J. Comp. Neurol. 505, 32– 45 (2007).

37. M. M. Díaz, M. Schlichting, K. C. Abruzzi, X. Long, M. Rosbash, Allatostatin-C/AstC-R2 Is a Novel Pathway to Modulate the Circadian Activity Pattern in Drosophila. Curr. Biol. CB. 29, 13–22.e3 (2019).

38. O. T. Shafer, C. Helfrich-Förster, S. C. P. Renn, P. H. Taghert, Reevaluation of Drosophila melanogaster’s Neuronal Circadian Pacemakers Reveals New Neuronal Classes. J. Comp. Neurol. 498, 180–193 (2006).

39. T. Goda, Y. Umezaki, F. Alwattari, H. W. Seo, F. N. Hamada, Neuropeptides PDF and DH31 hierarchically regulate free-running rhythmicity in Drosophila circadian locomotor activity. Sci. Rep. 9, 838 (2019).

40. C. Wegener, H. Herbert, J. Kahnt, M. Bender, J. M. Rhea, Deficiency of prohormone convertase dPC2 (AMONTILLADO) results in impaired production of bioactive neuropeptide hormones in Drosophila. J. Neurochem. 118, 581–595 (2011).

41. R. Lyutova, M. Selcho, M. Pfeuffer, D. Segebarth, J. Habenstein, A. Rohwedder, F. Frantzmann, C. Wegener, A. S. Thum, D. Pauls, Reward signaling in a recurrent circuit of dopaminergic neurons and peptidergic Kenyon cells. Nat. Commun. 10, 3097 (2019).

42. S. Dorkenwald, A. Matsliah, A. R. Sterling, P. Schlegel, S. Yu, C. E. McKellar, A. Lin, M. Costa, K. Eichler, Y. Yin, W. Silversmith, C. Schneider-Mizell, C. S. Jordan, D. Brittain, A. Halageri, K. Kuehner, O. Ogedengbe, R. Morey, J. Gager, K. Kruk, E. Perlman, R. Yang, D. Deutsch, D. Bland, M. Sorek, R. Lu, T. Macrina, K. Lee, J. A. Bae, S. Mu, B. Nehoran, E. Mitchell, S. Popovych, J. Wu, Z. Jia, M. Castro, N. Kemnitz, D. Ih, A. S. Bates, N. Eckstein, J. Funke, F. Collman, D. D. Bock, G. S. X. E. Jefferis, H. S. Seung, M. Murthy, the F. Consortium, Neuronal wiring diagram of an adult brain (2023), p. 2023.06.27.546656,, doi:10.1101/2023.06.27.546656.

43. Z. Zheng, J. S. Lauritzen, E. Perlman, C. G. Robinson, M. Nichols, D. Milkie, O. Torrens, J. Price, C. B. Fisher, N. Sharifi, S. A. Calle-Schuler, L. Kmecova, I. J. Ali, B. Karsh, E. T. Trautman, J. A. Bogovic, P. Hanslovsky, G. S. X. E. Jefferis, M. Kazhdan, K. Khairy, S. Saalfeld, R. D. Fetter, D. D. Bock, A Complete Electron Microscopy Volume of the Brain of Adult Drosophila melanogaster. Cell. 174, 730–743.e22 (2018).

44. J. Buhmann, A. Sheridan, C. Malin-Mayor, P. Schlegel, S. Gerhard, T. Kazimiers, R. Krause, T. M. Nguyen, L. Heinrich, W.-C. A. Lee, R. Wilson, S. Saalfeld, G. S. X. E. Jefferis, D. D. Bock, S. C. Turaga, M. Cook, J. Funke, Automatic detection of synaptic partners in a whole-brain Drosophila electron microscopy data set. Nat. Methods. 18, 771–774 (2021).

45. L. Heinrich, D. Bennett, D. Ackerman, W. Park, J. Bogovic, N. Eckstein, A. Petruncio, J. Clements, S. Pang, C. S. Xu, J. Funke, W. Korff, H. F. Hess, J. Lippincott-Schwartz, S. Saalfeld, A. V. Weigel, Whole-cell organelle segmentation in volume electron microscopy. Nature. 599, 141–146 (2021).

46. Y. Aso, K. Grübel, S. Busch, A. B. Friedrich, I. Siwanowicz, H. Tanimoto, The mushroom body of adult Drosophila characterized by GAL4 drivers. J. Neurogenet. 23, 156–172 (2009).

47. M. Fekete, J. M. van Ree, R. J. Niesink, D. de Wied, Disrupting circadian rhythms in rats induces retrograde amnesia. Physiol. Behav. 34, 883–887 (1985).

48. K. Cho, Chronic “jet lag” produces temporal lobe atrophy and spatial cognitive deficits. Nat. Neurosci. 4, 567–568 (2001).

49. F. A. J. L. Scheer, M. F. Hilton, C. S. Mantzoros, S. A. Shea, Adverse metabolic and cardiovascular consequences of circadian misalignment. Proc. Natl. Acad. Sci. U. S. A. 106, 4453–4458 (2009).

50. N. Takeda, K. Maemura, Circadian clock and vascular disease. Hypertens. Res. Off. J. Jpn. Soc. Hypertens. 33, 645–651 (2010).

51. A. H. Tsang, M. Astiz, M. Friedrichs, H. Oster, Endocrine regulation of circadian physiology. J. Endocrinol. 230, R1–R11 (2016).

52. A. Videnovic, A. S. Lazar, R. A. Barker, S. Overeem, ’The clocks that time us’—circadian rhythms in neurodegenerative disorders. Nat. Rev. Neurol. 10, 683–693 (2014).

53. P. Pereyra, H. O. De La Iglesia, H. Maldonado, Training-to-testing intervals different from 24 h impair habituation in the crab Chasmagnathus. Physiol. Behav. 59, 19–25 (1996).

54. M. R. Ralph, C. H. Ko, E. A. Antoniadis, P. Seco, F. Irani, C. Presta, R. J. McDonald, The significance of circadian phase for performance on a reward-based learning task in hamsters. Behav. Brain Res. 136, 179–184 (2002).

55. S. W. Cain, T. Chou, M. R. Ralph, Circadian modulation of performance on an aversion-based place learning task in hamsters. Behav. Brain Res. 150, 201–205 (2004).

56. F. A. Holloway, R. A. Wansley, Multiple retention deficits at periodic intervals after active and passive avoidance learning. Behav. Biol. 9, 1–14 (1973).

57. F. A. Holloway, R. Wansley, Multiphasic retention deficits at periodic intervals after passive-avoidance learning. Science. 180, 208–210 (1973).

58. N. S. Chouhan, R. Wolf, C. Helfrich-Förster, M. Heisenberg, Flies remember the time of day. Curr. Biol. CB. 25, 1619–1624 (2015).

59. N. S. Chouhan, R. Wolf, M. Heisenberg, Starvation promotes odor/feeding-time associations in flies. Learn. Mem. Cold Spring Harb. N. 24, 318–321 (2017).

60. D. Chaudhury, C. S. Colwell, Circadian modulation of learning and memory in fear-conditioned mice. Behav. Brain Res. 133, 95–108 (2002).

61. O. Rawashdeh, N. H. de Borsetti, G. Roman, G. M. Cahill, Melatonin suppresses nighttime memory formation in zebrafish. Science. 318, 1144–1146 (2007).

62. L. C. Lyons, C. L. Green, A. Eskin, Intermediate-Term Memory is Modulated by the Circadian Clock. J. Biol. Rhythms. 23, 538–542 (2008).

63. M. Kaneko, J. C. Hall, Neuroanatomy of cells expressing clock genes in Drosophila: transgenic manipulation of the period and timeless genes to mark the perikarya of circadian pacemaker neurons and their projections. J. Comp. Neurol. 422, 66–94 (2000).

64. T. Sakai, T. Tamura, T. Kitamoto, Y. Kidokoro, A clock gene, period, plays a key role in long-term memory formation in Drosophila. Proc. Natl. Acad. Sci. U. S. A. 101, 16058–16063 (2004).

65. J. H. Houl, W. Yu, S. M. Dudek, P. E. Hardin, Drosophila CLOCK is constitutively expressed in circadian oscillator and non-oscillator cells. J. Biol. Rhythms. 21, 93–103 (2006).

66. M. Heisenberg, Mushroom body memoir: from maps to models. Nat. Rev. Neurosci. 4, 266–275 (2003).

67. K. Eichler, F. Li, A. Litwin-Kumar, Y. Park, I. Andrade, C. M. Schneider-Mizell, T. Saumweber, A. Huser, C. Eschbach, B. Gerber, R. D. Fetter, J. W. Truman, C. E. Priebe, L. F. Abbott, A. S. Thum, M. Zlatic, A. Cardona, The complete connectome of a learning and memory centre in an insect brain. Nature. 548, 175–182 (2017).

68. P. Cognigni, J. Felsenberg, S. Waddell, Do the right thing: neural network mechanisms of memory formation, expression and update in Drosophila. Curr. Opin. Neurobiol. 49, 51–58 (2018).

69. F. Li, J. W. Lindsey, E. C. Marin, N. Otto, M. Dreher, G. Dempsey, I. Stark, A. S. Bates, M. W. Pleijzier, P. Schlegel, A. Nern, S.-Y. Takemura, N. Eckstein, T. Yang, A. Francis, A. Braun, R. Parekh, M. Costa, L. K. Scheffer, Y. Aso, G. S. Jefferis, L. F. Abbott, A. Litwin-Kumar, S. Waddell, G. M. Rubin, The connectome of the adult Drosophila mushroom body provides insights into function. eLife. 9, e62576 (2020).

70. G. Skofitsch, D. M. Jacobowitz, Calcitonin gene-related peptide: Detailed immunohistochemical distribution in the central nervous system. Peptides. 6, 721–745 (1985).

71. H. T. Park, S. Y. Baek, B. S. Kim, J. B. Kim, J. Jeong Kim, Calcitonin gene-related peptide-like immunoreactive (CGRPI) elements in the circadian system of the mouse: an immunohistochemistry combined with retrograde transport study. Brain Res. 629, 335–341 (1993).

72. Z. Kee, X. Kodji, S. D. Brain, The Role of Calcitonin Gene Related Peptide (CGRP) in Neurogenic Vasodilation and Its Cardioprotective Effects. Front. Physiol. 9 (2018) (available at https://www.frontiersin.org/articles/10.3389/fphys.2018.01249).

73. Y. Singh, G. Gupta, B. Shrivastava, R. Dahiya, J. Tiwari, M. Ashwathanarayana, R. K. Sharma, M. Agrawal, A. Mishra, K. Dua, Calcitonin gene-related peptide (CGRP): A novel target for Alzheimer’s disease. CNS Neurosci. Ther. 23, 457–461 (2017).

74. M. P. Fernández, J. Berni, M. F. Ceriani, Circadian remodeling of neuronal circuits involved in rhythmic behavior. PLoS Biol. 6, e69 (2008).

75. E. A. Gorostiza, A. Depetris-Chauvin, L. Frenkel, N. Pírez, M. F. Ceriani, Circadian pacemaker neurons change synaptic contacts across the day. Curr. Biol. CB. 24, 2161–2167 (2014).

76. B. J. Song, S. J. Sharp, D. Rogulja, Daily rewiring of a neural circuit generates a predictive model of environmental light. Sci. Adv. 7, eabe4284 (2021).

77. D. Owald, J. Felsenberg, C. B. Talbot, G. Das, E. Perisse, W. Huetteroth, S. Waddell, Activity of Defined Mushroom Body Output Neurons Underlies Learned Olfactory Behavior in Drosophila. Neuron. 86, 417–427 (2015).

78. E. Perisse, D. Owald, O. Barnstedt, C. B. Talbot, W. Huetteroth, S. Waddell, Aversive Learning and Appetitive Motivation Toggle Feed-Forward Inhibition in the Drosophila Mushroom Body. Neuron. 90, 1086–1099 (2016).

79. I. Cervantes-Sandoval, A. Phan, M. Chakraborty, R. L. Davis, Reciprocal synapses between mushroom body and dopamine neurons form a positive feedback loop required for learning. eLife. 6 (2017), doi:10.7554/eLife.23789.

80. N. Sgammeglia, S. G. Sprecher, Interplay between metabolic energy regulation and memory pathways in Drosophila. Trends Neurosci. 0 (2022), doi:10.1016/j.tins.2022.04.007.

81. M. J. Krashes, S. DasGupta, A. Vreede, B. White, J. D. Armstrong, S. Waddell, A neural circuit mechanism integrating motivational state with memory expression in Drosophila. Cell. 139, 416–427 (2009).

82. P.-Y. Plaçais, T. Preat, To favor survival under food shortage, the brain disables costly memory. Science. 339, 440–442 (2013).

83. D. Pauls, M. Selcho, J. Räderscheidt, K. M. Amatobi, A. Fekete, M. Krischke, C. Hermann-Luibl, A. G. Ozbek-Unal, N. Ehmann, P. M. Itskov, R. J. Kittel, C. Helfrich-Förster, R. P. Kühnlein, M. J. Mueller, C. Wegener, Endocrine signals fine-tune daily activity patterns in Drosophila. Curr. Biol. CB. 31, 4076–4087.e5 (2021).

84. K. Xu, X. Zheng, A. Sehgal, Regulation of feeding and metabolism by neuronal and peripheral clocks in Drosophila. Cell Metab. 8, 289–300 (2008).

85. T. Zars, M. Fischer, R. Schulz, M. Heisenberg, Localization of a short-term memory in Drosophila. Science. 288, 672–675 (2000).

86. M. Sekiguchi, K. Inoue, T. Yang, D.-G. Luo, T. Yoshii, A Catalog of GAL4 Drivers for Labeling and Manipulating Circadian Clock Neurons in Drosophila melanogaster. J. Biol. Rhythms. 35, 207–213 (2020).

87. Y. Shang, L. C. Griffith, M. Rosbash, Light-arousal and circadian photoreception circuits intersect at the large PDF cells of the Drosophila brain. Proc. Natl. Acad. Sci. U. S. A. 105, 19587– 19594 (2008).

88. J. M. Rhea, C. Wegener, M. Bender, The proprotein convertase encoded by amontillado (amon) is required in Drosophila corpora cardiaca endocrine cells producing the glucose regulatory hormone AKH. PLoS Genet. 6, e1000967 (2010).

89. M. D. Gordon, K. Scott, Motor control in a Drosophila taste circuit. Neuron. 61, 373–384 (2009).

90. B. Schmid, C. Helfrich-Förster, T. Yoshii, A new ImageJ plug-in “ActogramJ” for chronobiological analyses. J. Biol. Rhythms. 26, 464–467 (2011).

