## Supplementary Figures and Table for "Neuronal correlates of time integration into memories"

### Supplementary Material

**Frantzmann et al.**

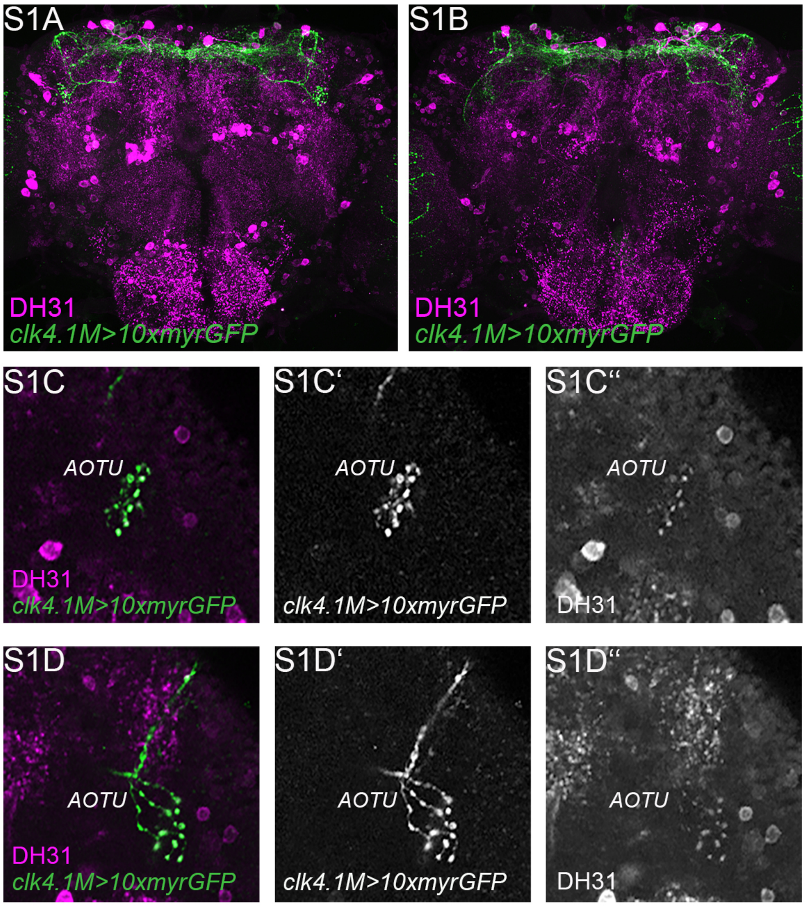


***Supplement Figure S1: Overlap of DN1p neurons and DH31 antibody staining. (A,B)*** *Anterior (A) and posterior (B) view of whole mount projections of brains visualizing clk4.1M-Gal4-positive neurons (green) and DH31-positive cells (magenta).* ***(C-D"')*** *Only type B DN1p neurons innervate the anterior optic tubercle (AOTU). All DN1p terminals in the AOTU are DH31-positive.[AOTU: Anterior optic tubercle; DH31: Diuretic hormone 31; DN1p: Posterior dorsal neuron]*

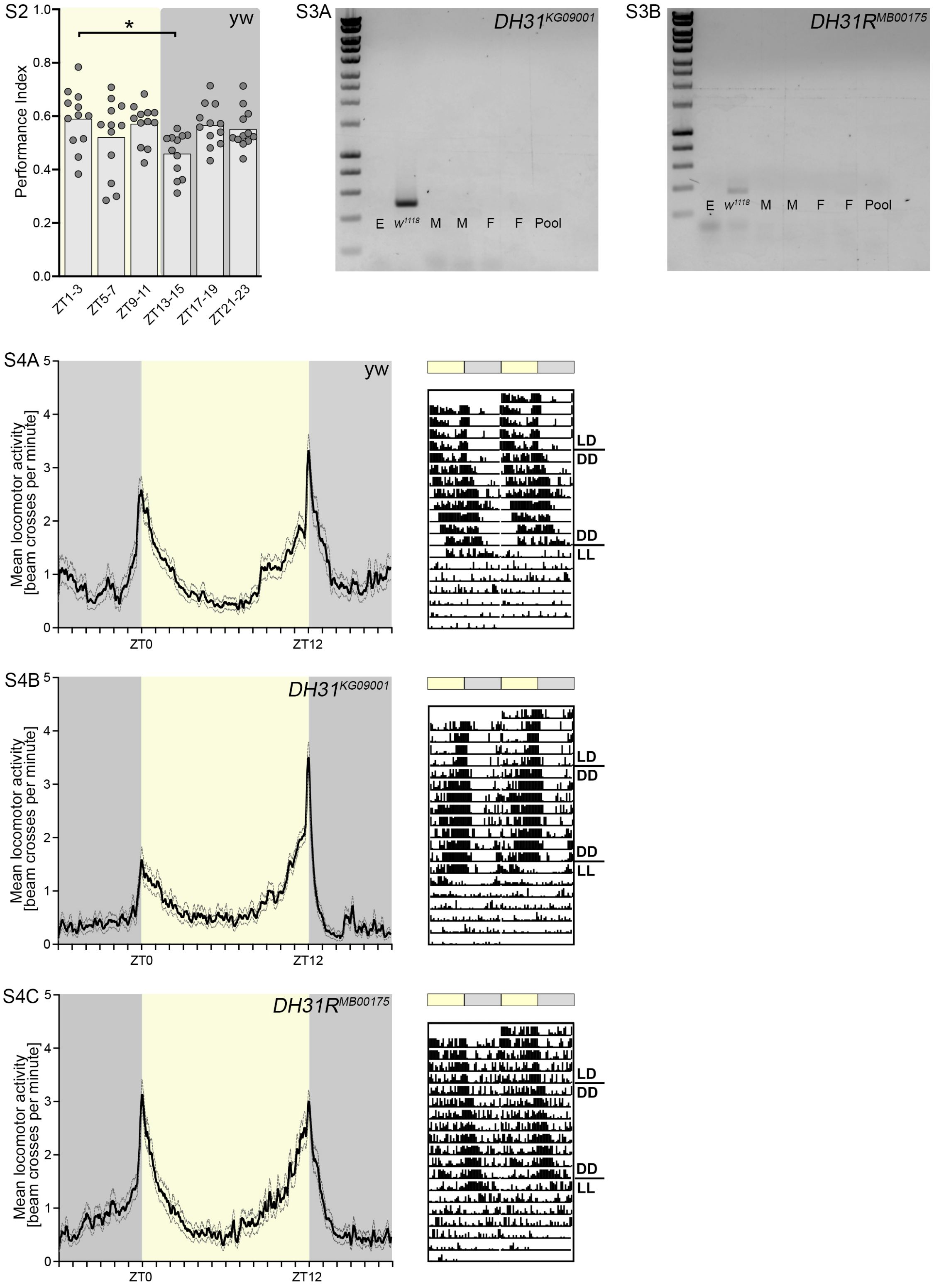


***Supplement Figure S2-S4:*** ***(S2)*** *Memory performance for time slots during the LD cycle (day: ZT1-3, ZT5-7 and ZT9-11; night: ZT13-15, ZT17-19 and ZT21-23) are shown for control flies (grey). Only mild fluctuations are visible throughout 24h.* ***(S3)*** *Genotyping of* $DH31^{KG09001}$ *and* $DH31R^{MB00175}$ *mutant lines.* ***(S4)*** *Locomotor activity of DH31* ***(S4B)*** *and DH31R flies* ***(S4C)*** *compared to yw control flies (S4A). Activity profiles and exemplary actograms indicate that flies lacking DH31 signalling show an overall rhythmical behaviour under LD conditions, indicating that the lack of DH31 does not fully impair clock function. In line, also under constant darkness (DD) flies are rhythmic, while constant light (LL) completely impairs clock function. [DH31: Diuretic hormone 31; DH31R: DH31 receptor; E: Empty; F: Female; M: Male; ZT: Zeitgeber time]*

| FlyWire segment ID | Contributor (min 10% of total edits) | Annotation |
|---|---|---|
| **720575940613413791** | Mareike Selcho | Mareike Selcho |
| **720575940619067259** | Mareike Selcho, Doug Bland | Mareike Selcho |
| **720575940631973089** | Imaan Tamimi, Mareike Selcho | Mareike Selcho |
| **720575940642237344** | Mareike Selcho | Mareike Selcho |
| **720575940620561713** | Mareike Selcho | Mareike Selcho |
| **720575940622398836** | Mareike Selcho | Mareike Selcho |
| **720575940628257874** | Darrel Jay Akiatan, Joshua Bañez, Mareike Selcho | Nils Reinhard, Dustin Garner |
| **720575940628692040** | Mareike Selcho, Zeba Vohra, Zairene Lenizo | Mareike Selcho |
| **720575940644609440** | Shirleyjoy Serona, Doug Bland | Nils Reinhard |
| **720575940645975063** | Dustin Garner, Mareike Selcho | Nils Reinhard, Dustin Garner |
| **720575940625455134** | Arti Yadav, Anjali Pandey, A. Javier | Mareike Selcho |
| **720575940614792086** | J. Anthony Ocho, Arti Yadav, A. Javier | Wolf Huetteroth |
| **720575940630705745** | Mareike Selcho, Arti Yadav, A. Javier, Monika Patel, Rashmita Rana | Mareike Selcho |
| **720575940621576063** | Michelle Pantujan, Itisha Joshi, A. Javier | Mareike Selcho |
| **720575940641765773** | Arti Yadav, Katharina Eichler, A. Javier, Irene Salgarella | Mareike Selcho, Volker Hartenstein, Alexander Bates |
| **720575940621948554** | Doug Bland, Mareike Selcho, A. Javier, Arti Yadav | Mareike Selcho |
| **720575940622225043** | Mendell Lopez, A. Javier, Arti Yadav | Mareike Selcho |
| **720575940618400459** | Claire McKellar, Anjali Pandey, Arti Yadav | Mareike Selcho |
| **720575940604505278** | Dharini Sapkal, Zairene Lenizo, Arti Yadav | Wolf Huetteroth |
| **720575940635471608** | Anjali Pandey, A. Javier, Bhargavi Parmar | Mareike Selcho |
| **720575940613757224** | Dhwani Patel, Arti Yadav, Chitra Nair, Anjali Pandey, Bhargavi Parmar | Wolf Huetteroth |
| **720575940610872418** | Doug Bland, A. Javier, Dharini Sapkal | Mareike Selcho |
| **720575940618756987** | Doug Bland, Monika Patel, Dharini Sapkal | Wolf Huetteroth |
| **720575940626560601** | Arti Yadav, A. Javier, Dhwani Patel | Wolf Huetteroth |
| **720575940632985495** | Anjali Pandey, A. Javier, Irene Salgarella | Mareike Selcho, Volker Hartenstein, Alexander Bates |
| **720575940625607946** | Arti Yadav, A. Javier, Irene Salgarella | Mareike Selcho, Volker Hartenstein, Alexander Bates |
| **720575940624882702** | A. Javier, Anjali Pandey, Irene Salgarella | Mareike Selcho, Volker Hartenstein, Alexander Bates |
| **720575940603687084** | A. Javier, Anjali Pandey, Irene Salgarella, Monika Patel | Mareike Selcho, Volker Hartenstein, Alexander Bates |
| **720575940636571310** | Doug Bland, Dhara Kakadiya, Irene Salgarella | Mareike Selcho, Volker Hartenstein, Alexander Bates |
| **720575940614967007** | Albert Lin, Arti Yadav, Kaushik Parmar | Wolf Huetteroth |
| **720575940615395414** | Arti Yadav, Greg Jefferis, Kyle Patrick Willie | Wolf Huetteroth |
| **720575940616750363** | Dharini Sapkal, Anjali Pandey, Mareike Selcho | Mareike Selcho |
| **720575940615830877** | Arti Yadav, Shirleyjoy Serona, Mareike Selcho | Mareike Selcho, Wolf Huetteroth |
| **720575940633550305** | Doug Bland, Anjali Pandey, Michelle Pantujan | Mareike Selcho |
| **720575940637832255** | Anjali Pandey, A. Javier, Monika Patel, Mendell Lopez | Mareike Selcho |
| **720575940604435046** | Anjali Pandey, A. Javier, Monika Patel | Mareike Selcho, Volker Hartenstein, Alexander Bates |
| **720575940619942320** | Kyle Patrick Willie, Arti Yadav, Monika Patel | Wolf Huetteroth |
| **720575940625991524** | Anjali Pandey, Irene Salgarella, Monika Patel, Itisha Joshi | Mareike Selcho, Volker Hartenstein, Alexander Bates |
| **720575940616560541** | Dharini Sapkal, James Hebditch, Philipp Schlegel, Arti Yadav | Mareike Selcho |
| **720575940615904145** | Nash Hadjerol, Anjali Pandey, Rashmita Rana | Wolf Huetteroth |
| **720575940637787226** | Anjali Pandey, Rashmita Rana, Yashvi Patel, Kaushik Parmar | Mareike Selcho |
| **720575940620810316** | Arti Yadav, A. Javier | Mareike Selcho |
| **720575940614842741** | Arti Yadav, A. Javier | Mareike Selcho, Volker Hartenstein, Alexander Bates |
| **720575940617143966** | Doug Bland, A. Javier | Mareike Selcho, Volker Hartenstein, Alexander Bates |
| **720575940625605257** | Mareike Selcho, Anjali Pandey | Mareike Selcho |
| **720575940628431042** | Mareike Selcho, Anjali Pandey | Mareike Selcho |
| **720575940628411002** | Anjali Pandey, Arti Yadav | Mareike Selcho |
| **720575940631843980** | Dhara Kakadiya, Arti Yadav | Mareike Selcho |
| **720575940617687330** | Dhwani Patel, Arti Yadav | Mareike Selcho |
| **720575940607168523** | Doug Bland, Arti Yadav | Wolf Huetteroth |
| **720575940606851714** | Nash Hadjerol, Bhargavi Parmar | Wolf Huetteroth |
| **720575940631093007** | Alisa Poh, Chitra Nair | Mareike Selcho |
| **720575940625903626** | Arti Yadav, Dharini Sapkal | Mareike Selcho |

| | | |
|---|---|---|
| **720575940613784574** | Anjali Pandey, Dhwani Patel | Wolf Huetteroth |
| **720575940609676100** | Arti Yadav, Dhwani Patel | Wolf Huetteroth |
| **720575940645975063** | Dustin Garner, Mareike Selcho | Dustin Garner, Nils Reinhard |
| **720575940614562098** | Anjali Pandey, Markus Pleijzier | Wolf Huetteroth |
| **720575940626838513** | Arti Yadav, Monika Patel | Mareike Selcho |
| **720575940630263619** | Arti Yadav, Shirleyjoy Serona | Mareike Selcho |
| **720575940618626102** | Anjali Pandey, Yashvi Patel | Mareike Selcho |
| **720575940623260540** | Austin T Burke, Zairene Lenizo | Mareike Selcho |
| **720575940627805928** | Arti Yadav, Zeba Vohra | Mareike Selcho |
| **720575940626580784** | Mareike Selcho, Zeba Vohra | Wolf Huetteroth, Mareike Selcho |
| **720575940624153608** | Arti Yadav | Mareike Selcho |
| **720575940611213774** | Arti Yadav | Mareike Selcho |
| **720575940620467836** | Arti Yadav | Mareike Selcho |
| **720575940627048522** | J. Dolorosa | Mareike Selcho |
| **720575940617175413** | Mareike Selcho | Mareike Selcho |

**Table S1:** Overview of proofreading and annotations of the cells shown in this publication.
